## Supplementary figures and images for "Phytotoxin synthesis genes and type III effector genes of *Pseudomonas syringae* pv. *actinidiae* biovar 6 are regulated by culture conditions"

### Supplemental Figure S1.pdf

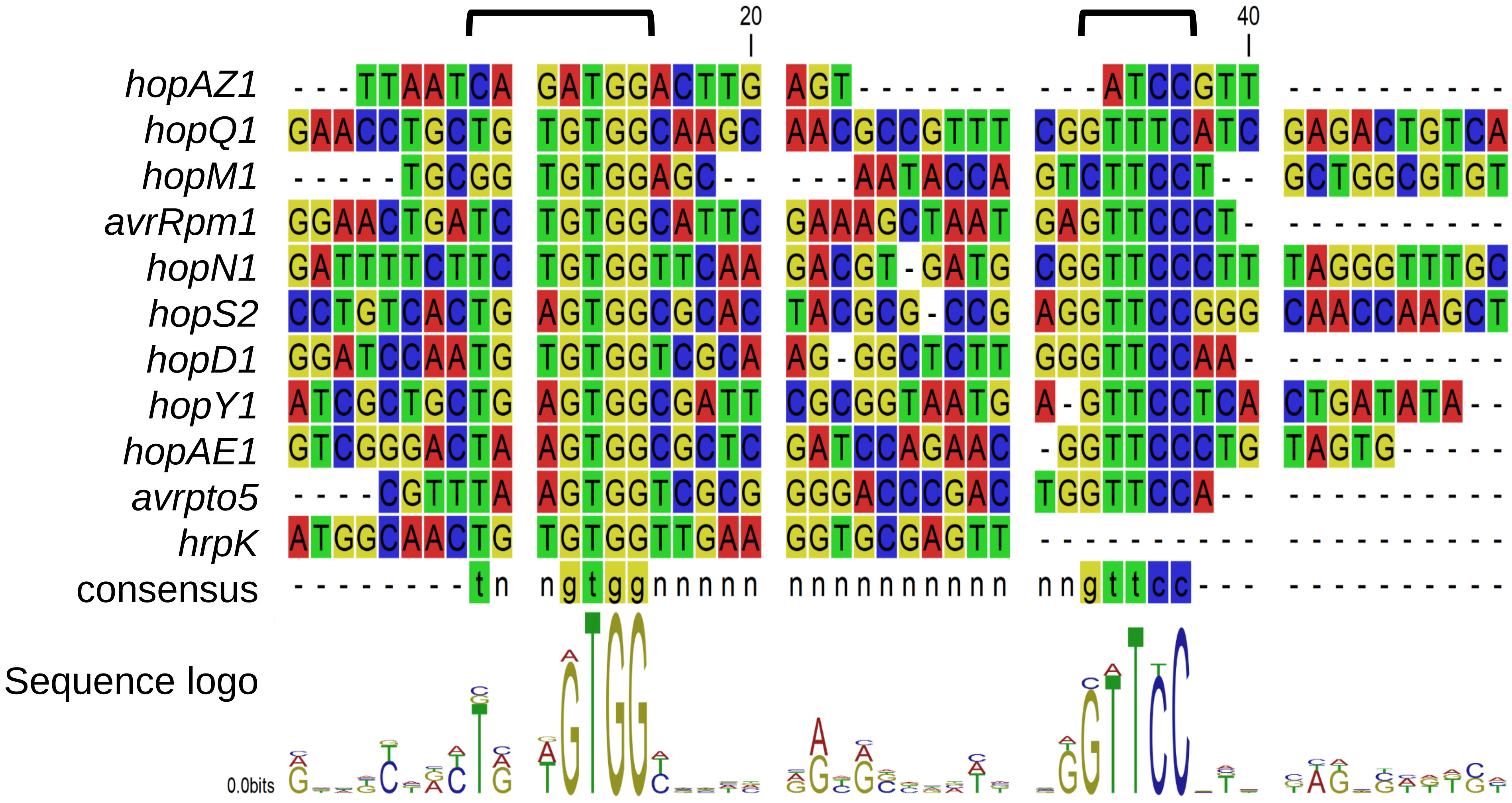
